# Investigating the mechanisms underlying saccade generation in the frontal eye fields using multi-site microstimulation

**DOI:** 10.64898/2025.12.29.696832

**Authors:** Richard Johnston, Roma O. Konecky, Husam A. Katnani, Neeraj J. Gandhi, Matthew A. Smith

## Abstract

The frontal eye field (FEF), a region of frontal cortex, has long been associated with the cortical control of eye movements. Classically, saccades can be reliably evoked by delivering low-intensity electrical microstimulation to the FEF. Although this makes clear the importance of FEF in the descending control of eye movements, the way in which population activity in the FEF is integrated by downstream regions to generate a motor command remains a mystery. To probe these mechanisms, we used a 16-channel microelectrode array to deliver microstimulation to the FEF of two awake, behaving monkeys. First, we found that larger current intensities were required to evoke changes in saccade direction relative to saccade amplitude when single-site saccades were evoked by stimulating a single contact on the array. Second, when stimulating two contacts simultaneously to investigate how population activity in the FEF is read out, a new polar average model more accurately predicted the amplitude and direction of dual-site saccades than traditional vector sum and vector average models. Using preexisting data from the superior colliculus (SC), we found that although the polar average model was more accurate at predicting saccade amplitude in the SC, it was no more accurate than traditional models at predicting saccade direction. Finally, when stimulating two contacts in FEF simultaneously with unequal current intensities, model accuracy depended on the amplitude of the saccades evoked by stimulating each individual site alone, suggesting that the brain may flexibly combine amplitude and direction information from the FEF to generate saccadic plans.

## Introduction

The frontal eye field (FEF) has long been associated with the generation of saccadic eye movements (Grosbras et al. 2005; Paus 1996; Schall 2015; Tehovnik et al. 2000; Vernet et al. 2014). This region, which resides in the bank of the arcuate sulcus, contains neurons that can be divided into three traditional classes: visual neurons that respond when stimuli fall within their receptive field (RF), motor neurons that respond when an eye movement is made into their RF and visuomotor neurons that exhibit visual- and saccade-related responses (Bruce and Goldberg 1985; Khanna et al. 2019; Lawrence and Snyder 2009; Lowe and Schall 2018). Saccade amplitude is topographically organized in the FEF; small saccades are encoded by neurons in the ventrolateral portion and large saccades by neurons in the dorsomedial portion (Babapoor-Farrokhran et al. 2013; Bruce et al. 1985; Robinson and Fuchs 1969; Selvanayagam et al. 2019; Stanton et al. 1988a). In contrast, saccade direction is less systematically organized in the FEF, especially when compared to the superior colliculus (SC). The SC is interconnected with the FEF (Crapse and Sommer 2009; Ferraina et al. 2002; Fries 1984; Komatsu and Suzuki 1985; Paré and Wurtz 2001; Sommer and Wurtz 2000; Stanton et al. 1988b) and also contains neurons with visual, motor and visuomotor responses (Bourrelly et al. 2023; Dorris et al. 1997; Gandhi and Katnani 2011; Goldberg and Wurtz 1972; Massot et al. 2019; Wurtz and Albano 1980; Wurtz and Goldberg 1972). Like the FEF, saccade amplitude is well organized in the SC; small saccades are encoded by neurons in the rostral portion and large saccades by neurons in the ventral portion (Robinson 1972; Schiller and Stryker 1972; Van Opstal et al. 1990). However, in contrast to the FEF, saccade direction is also organized topographically in the SC; upward saccades are encoded by neurons in the medial portion and downward saccades by neurons in the ventral portion (Bruce et al. 1985; Dias and Segraves 1999; Robinson and Fuchs 1969). Consequently, neurons in the SC are thought to form a retinotopically organized map in which neighboring neurons exhibit similar preferences for saccade amplitude and direction, whereas the topographical organization of the FEF is more patchy and less continuous, particularly with respect to saccade direction.

Although the FEF and SC differ in topographical organization, one common characteristic of these two regions is that saccades can be evoked via intracortical microstimulation at relatively low current intensities (Bruce et al. 1985; Goldberg et al. 1986; Katnani and Gandhi 2011, 2012; Paré et al. 1994; Robinson 1972; Robinson and Fuchs 1969; Schall 1991; Schiller and Stryker 1972; Stanford et al. 1996; Tehovnik 1996; Tehovnik and Lee 1993; Tehovnik and Sommer 1997; Van Opstal et al. 1990). In the SC, the amplitude of saccades evoked by single-site microstimulation (i.e., stimulation at a single location, typically performed by advancing an electrode into the SC) depends on the stimulation parameters used, such that saccade amplitude increases with current intensity and frequency (Katnani et al. 2012; Katnani and Gandhi 2012; Paré et al. 1994). However, it is important to note that the maximum amplitude of a saccade can be reached using non-optimal parameters if train duration is increased, highlighting the importance of controlling for pulse number in addition to current intensity and frequency (Stanford et al. 1996). In the FEF, the probability of evoking single-site saccades also varies with stimulation parameters (Murphey and Maunsell 2008; Robinson and Fuchs 1969; Tehovnik 1996; Tehovnik and Lee 1993; Tehovnik and Sommer 1997) but less is known about how these parameters impact saccade amplitude, direction and velocity. Given the differences in topographical organization between the FEF and SC (particularly the less orderly encoding of saccade direction in the FEF), it is possible that the effects of current intensity and frequency on the amplitude and direction of single-site saccades may differ between these two regions.

Another question that has received less attention in the FEF than the SC is how population activity is interpreted (or “read-out”) by downstream regions to generate a saccadic eye movement. In the SC, two main decoding algorithms have been proposed: vector summation and vector averaging (Brecht et al. 2004; Katnani et al. 2012; Lee et al. 1988; van Opstal and Goossens 2008; Van Gisbergen et al. 1987). Although these models make identical predictions with respect to saccade direction, the vector sum model tends to predict larger amplitude saccades relative to the vector average model. To study how saccade generation signals are summed in the SC, Brecht et al. (2004) stimulated two sites with equal current intensities. The endpoints of the resulting dual-site saccades were then compared to the predictions of the vector sum and vector average models, which were computed using the endpoints of saccades evoked by stimulating each individual site alone. Results showed that the amplitude and direction of the dual-site saccades closely followed the predictions of the vector average model when the timing of microstimulation across the two sites was synchronous. In contrast, the dual-site saccades followed the predictions of the vector sum model when the timing of microstimulation across the two individual sites was asynchronous. These findings can be interpreted within the framework of the vector summation with saturation model (Groh 2001), which predicts that saccade vectors are summed at relatively low activity levels and averaged at relatively high activity levels. Although there is some support for this model (Goossens and Van Opstal 2006; Groh 2001), Katnani and Gandhi (2012) found that a weighted vector average model performed better than the vector summation model both when dual-site saccades were evoked using suprathreshold microstimulation parameters with reduced parameters that produced saccades with a nonoptimal amplitude and direction. Thus, the vector average model appears to provide a better prediction of saccades evoked by dual-site microstimulation in the SC than the vector sum model and the vector summation with saturation model. However, given the lack of studies that have performed dual-site microstimulation in the FEF, it is unclear if this computation generalizes across these two oculomotor regions, or whether it is specific to population decoding in the SC.

The overall goal of this study was to investigate how saccades are generated in the FEF using single- and dual-site microstimulation. Firstly, although previous work has studied how microstimulation parameters impact the probability of evoking single-site saccades in the FEF (Murphey and Maunsell 2008; Robinson and Fuchs 1969; Tehovnik 1996; Tehovnik and Lee 1993; Tehovnik and Sommer 1997), much less is known about how these parameters affect saccade amplitude, direction and velocity. To address this question in the present study, we applied microstimulation to single sites in the FEF of non-human primates (NHPs) at different current intensities and measured changes in these three metrics as well as the percentage of evoked saccades. Secondly, we investigated how population activity in the FEF is read-out by stimulating two sites in the SC simultaneously with equal current intensities. We tested the predictions of the traditional vector sum and vector average models, as well as a new polar average model in which amplitude and direction are represented separately. Thirdly, we leveraged preexisting dual-site stimulation data in the SC (Katnani and Gandhi 2011) to determine if a similar computational model is used to decode population activity in the SC and FEF. Given that saccade direction is less organized in the FEF relative to the SC, it is possible that a different computation may be used to decode population activity. Finally, after assessing whether the computation used to decode population activity in the FEF generalizes to the SC, we tested the predictions of the three models when dual-site microstimulation was performed with unequal current intensities across the two sites.

## Methods

### Subjects

Two adult rhesus macaque monkeys (*Macaca mulatta*) were used in this study. Surgical procedures to chronically implant a titanium head post (to immobilize the subjects’ heads during experiments) and recording chamber were performed under isoflurane anesthesia, as described previously (Mayo et al. 2015). Opiate analgesics were used to minimize pain and discomfort during the perioperative period. All experimental procedures were approved by the Institutional Animal Care and Use Committee of the University of Pittsburgh and were performed in accordance with the United States National Research Council’s Guide for the Care and Use of Laboratory Animals.

### Electrophysiology

Recordings were made with a 16-channel microelectrode array (U-Probe, Plexon, Dallas, TX) with contacts spaced 150 µm apart using a Ripple Grapevine system (Salt Lake City, Utah). Microstimulation was performed with Ripple dual stimulation-recording headstages (350 Hz biphasic, 0.25 μS pulse width, 15 – 150 μA). The probe was advanced into the FEF using an NIH-designed mechanical microdrive (Laboratory for Sensorimotor Research, Bethesda, MD) through a plastic grid with 1 mm spacing that was placed inside the recording chamber (Crist Instruments, Hagerstown, MD). We placed the recording chamber at specific coordinates to center on FEF (stereotaxic coordinates: 25 mm anterior, 20 mm lateral) and in the implantation surgery attempted to visualize the arcuate sulcus through the intact dura to guide our recording locations. We then confirmed the location of FEF in two ways. First, we recorded extracellular activity from each channel on the array while the monkeys performed a memory-guided saccade task (Khanna et al. 2020; Mayo et al. 2015). Waveform segments crossing a threshold (set as a multiple of the root mean square noise on each channel) were digitized (30 KHz) and stored for offline processing, which involved plotting the peristimulus time histogram for each channel and assessing changes in spiking activity during the visual, delay and saccade epochs of the task. Second, we applied microstimulation at single sites along the array and identified grid holes where fixed-vector saccades could be reliably evoked (>50% of the time) using low current intensities (≤50 μA for 70 ms).

### Visual stimuli and experimental task

The stimuli for the fixation task were generated using a combination of custom software written in MATLAB (The MathWorks) and Psychophysics Toolbox extensions (Brainard, 1997; Pelli, 1997; Kleiner et al., 2007). They were displayed on a CRT monitor (resolution = 1024 X 768 pixels; refresh rate = 100Hz), which was viewed at a distance of 36cm and gamma-corrected to linearize the relationship between input voltage and output luminance using a photometer and look-up tables. Stimulus timing and alignment was confirmed using a photodiode. To initiate a trial, the monkey fixated on a central point (diameter = 0.3°) (Figure 1A). After a randomized fixation period spanning 200-600 ms, the fixation point was extinguished for 50 ms. Microstimulation was then applied for 200 ms at either (1) a single site along the array or (2) two sites simultaneously. After the stimulation was complete, the monkey received a liquid reward regardless of whether or not its eyes left the invisible fixation window (diameter = 1.8°). After the completion of a trial, the monkey had to wait 1000 ms before initiating the next trial. Eye position was monitored using an infrared eye tracker (EyeLink 1000, SR Research, Mississauga, Canada).

**Figure 1.**
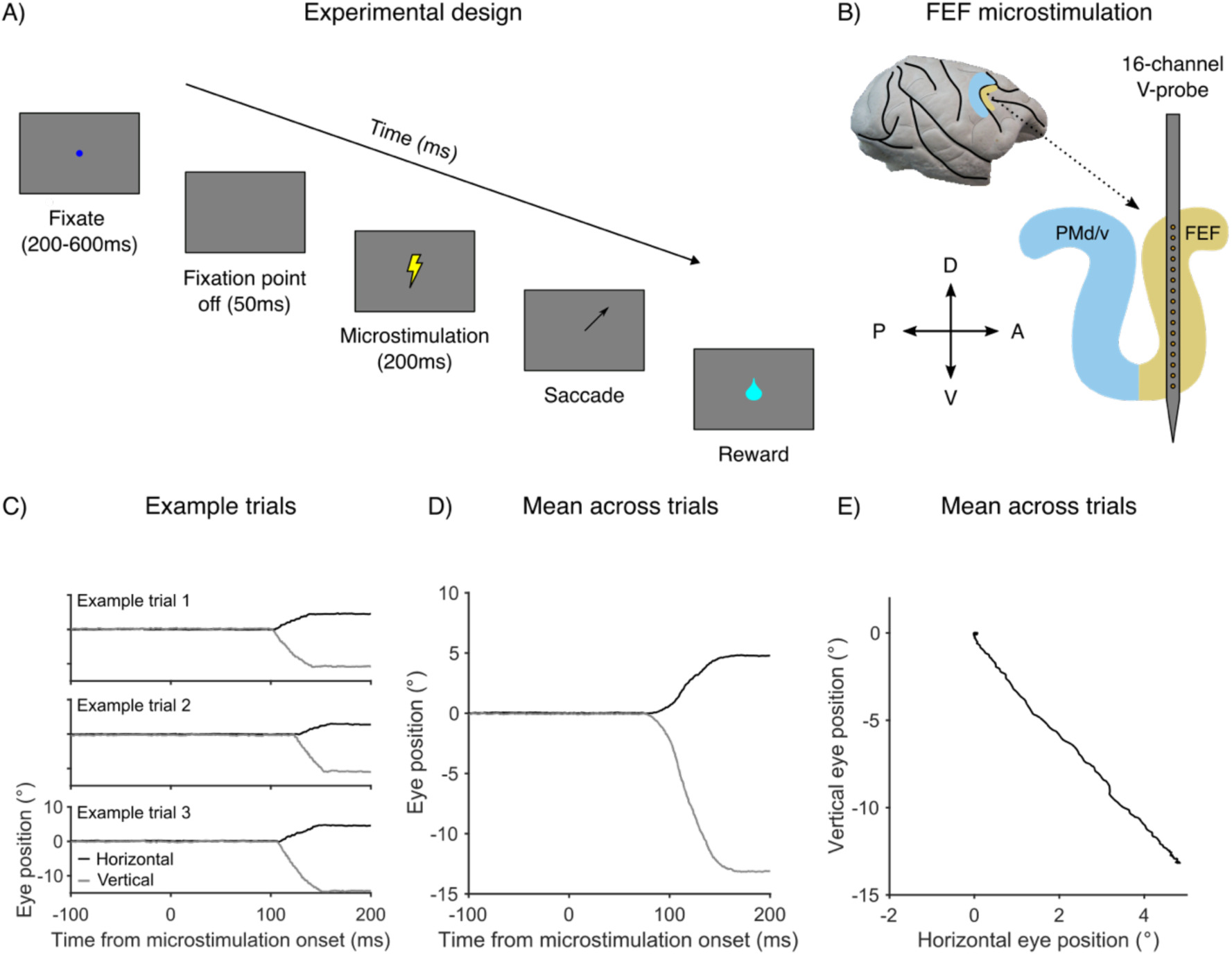
Single-site microstimulation in the FEF. (A) Schematic of the fixation task during which microstimulation was applied to evoke a saccadic eye movement. (B) Microstimulation was applied at a single contact on a 16-channel linear microelectrode array that was lowered into the FEF through a chronically implanted recording chamber. (C) Horizontal and vertical eye position in 1D space for three individual trials. (D) Mean horizontal and vertical eye position across trials for the same current intensity. (E) The same mean saccade vector as (D) but this time plotted in two-dimensional (2D) eye position space to better visualize the direction of the evoked saccade.

### Saccade detection

We wanted to ensure that our method for detecting saccades evoked by microstimulation was as sensitive as possible to small changes in eye position at low current intensities. Therefore, we utilized an approach that is commonly used to detect fixational saccades (Engbert and Kliegl 2003). Saccades were defined as eye movements that exceeded a velocity threshold of six times the standard deviation of the median velocity across the entire trial for at least 6 ms. When detecting fixational saccades, an upper bound on saccade amplitude is often imposed to exclude large eye movements. Here, we did not apply an amplitude limit, because neurons were targeted in the ventrolateral and dorsomedial portions of the FEF, which encode for small- and large-amplitude saccades, respectively. To ensure that the microstimulation train duration (fixed at 200 ms) was always long enough to allow for completion of the stimulation-evoked movement, we excluded trials in which the saccade did not drop below the threshold before the end of the stimulation period.

## Results

### Single-site microstimulation in the FEF

To study the effects of single-site microstimulation in the FEF, we trained two monkeys to perform a fixation task (Figure 1A). After a brief period of fixation on a central fixation point (200 – 600 ms), microstimulation was applied for 200 ms (350 Hz biphasic, 0.25 μS pulse width, 15 – 150 μA) at a single site along a 16-channel linear microelectrode array that was lowered into the FEF through a chronically implanted recording chamber (Figure 1B). If the microstimulation current intensity was sufficiently strong, an eye movement was evoked with a specific amplitude and direction. This can be seen in Figure 1C, which plots horizontal and vertical eye position for three individual trials in one-dimensional (1D) eye position space. Next, the mean saccade vector was computed by averaging across trials with the same current intensity (Figure 1D). Figure 1E shows the same mean saccade vector plotted in two-dimensional (2D) eye position space to better visualize the direction of the evoked saccade.

### Impact of current intensity on saccades evoked by single-site stimulation in the FEF

First, we investigated the impact of microstimulation current intensity on saccades evoked by single-site stimulation in the FEF. We focused on four metrics that have been used to assess the impact of stimulation parameters on single-site saccades: saccade amplitude, saccade direction, saccade velocity and the percentage of evoked saccades. In many example sites such as those shown in Figure 2A, we found that the amplitude of the mean saccade vector increased with microstimulation current intensity before reaching a plateau. A similar relationship was found for saccade direction. This was assessed by computing the absolute difference in direction (“theta”) between each individual mean saccade vector and the mean saccade vector evoked by the lowest amount of current. As shown in Figure 2B, the absolute theta difference increased with current intensity before reaching a plateau. A similar relationship was found for saccade velocity (Figure 2C) and the percentage of evoked saccades (Figure 2D).

**Figure 2.**
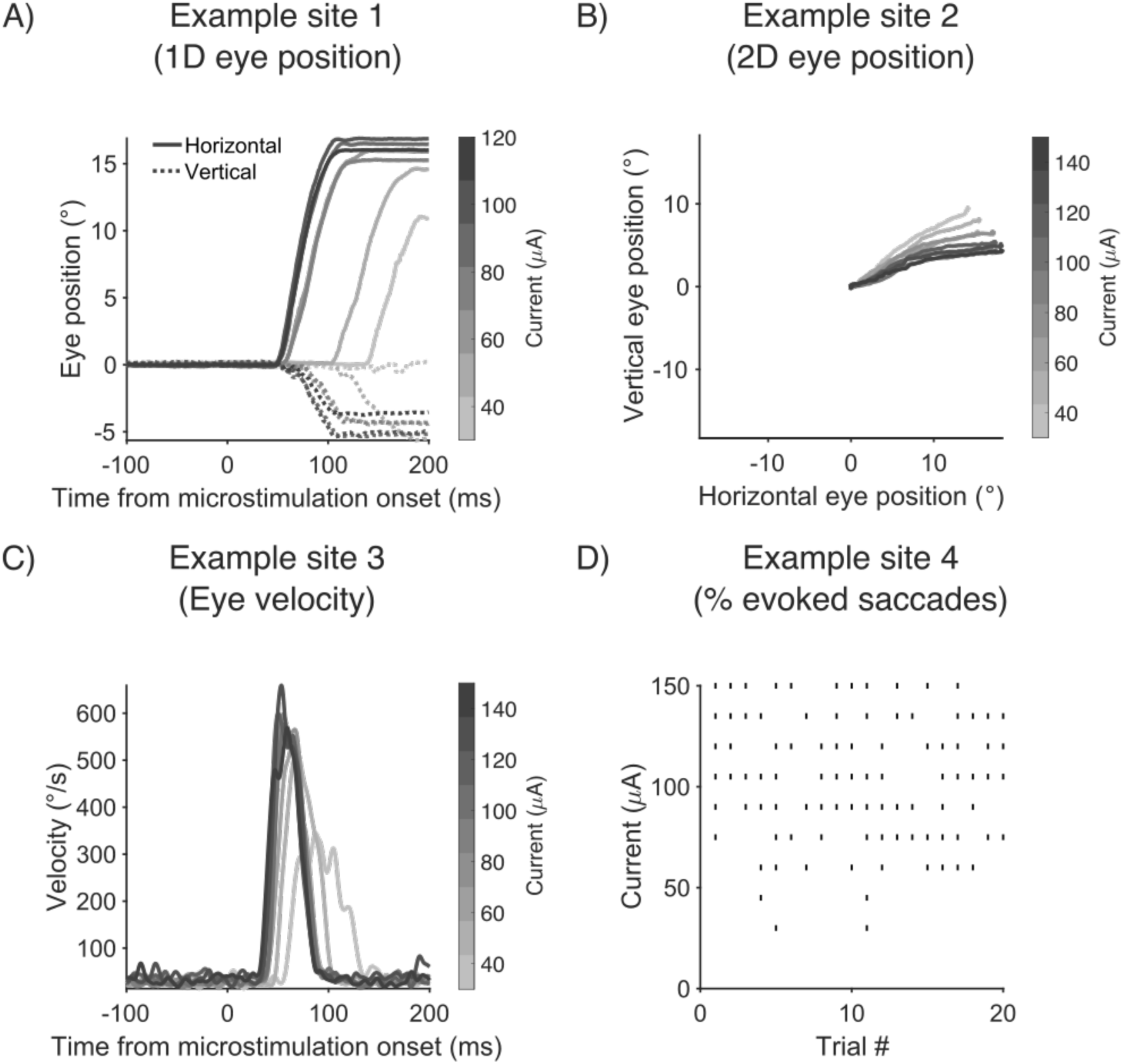
Impact of current intensity on saccades evoked by single-site stimulation in the FEF. (A) Mean horizontal and vertical eye position plotted in 1D space for a single site as a function of current intensity. (B) Mean horizontal and vertical eye position for a single site plotted in 2D space. Microstimulation current intensity is indicated by the color bar. (C) Mean saccade velocity for a single site. Microstimulation current intensity is indicated by the color bar. (D) Raster plot showing the number of evoked saccades across trials at each current level.

Next, we characterized the impact of microstimulation current intensity on saccades evoked by single-site stimulation by fitting a Naka-Rushton function to the data for each individual site (Peirce 2007). This allowed us to quantify the current intensity that was needed to evoke a 50% change in each metric relative to the saccade evoked at maximum current intensity (termed the “c50”). For the example site shown in Figure 2A, the c50 for saccade amplitude was 26.07 µA (Figure 3A), while for the site shown in Figure 2B, the c50 for saccade direction was 58.94 µA (Figure 3B). For the site shown in Figure 2C, the c50 for saccade velocity was 26.07 µA (Figure 3C), while for the site shown in Figure 2D, a current intensity of 54.79 µA was needed to evoke a 50% decrease from the maximum percentage of evoked saccades (Figure 3D).

**Figure 3.**
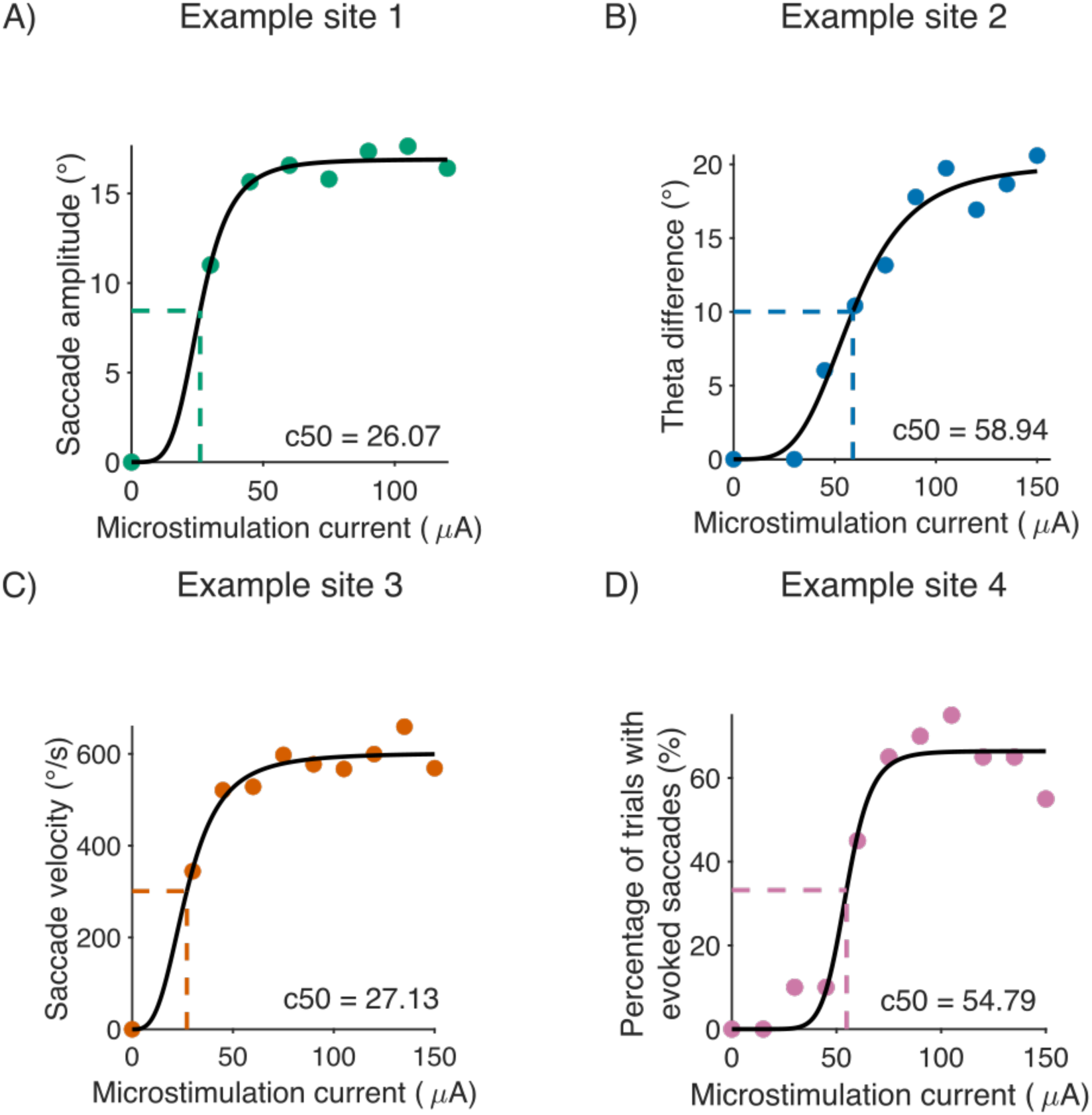
Impact of current intensity on saccades evoked by single-site stimulation in the FEF. (A) Scatterplot showing the relationship between current intensity and saccade amplitude for the site shown in Figure 2A. (B) Scatter plot showing the relationship between current intensity and theta difference (defined as the absolute difference in direction between each individual mean saccade vector and the mean saccade vector evoked by the lowest amount of current) for the site shown in Figure 2B. (C) Scatter plot showing the relationship between current intensity and saccade velocity for the site shown in Figure 2C. (D) Scatter plot showing the relationship between current intensity and the percentage of trials with evoked saccades for the site shown in Figure 2D. In each panel, a Naka-Rushton function was fit to the data to quantify the current needed to evoke a 50% change relative to the saccade evoked at maximum current.

To determine if similar results were found across all single sites, we generated a histogram of c50 values for each saccade metric. We limited the sites included in this analysis to ones where the *r^2^* of the Naka-Rushton function was greater than 0.7. The mean current needed to evoke a 50% change in saccade amplitude relative to the saccade evoked at maximum current was 21.37 µA (Figure 4A). In contrast, the mean c50 for saccade direction was twice as large (45.17 µA, Figure 4B), consistent with what was observed across example sites (Figure 3A and Figure 3B), and demonstrating that a significantly greater amount of current is needed to evoke a 50% change in saccade direction relative to saccade amplitude (*t* (26) = -6.21, *p* < 0.001, paired-sample *t*-test). For saccade velocity, the mean c50 was 21.52 µA (Figure 4C), whereas a mean current intensity of 40.42 µA was needed to evoke a 50% decrease from the maximum percentage of evoked saccades (Figure 4D).

**Figure 4.**
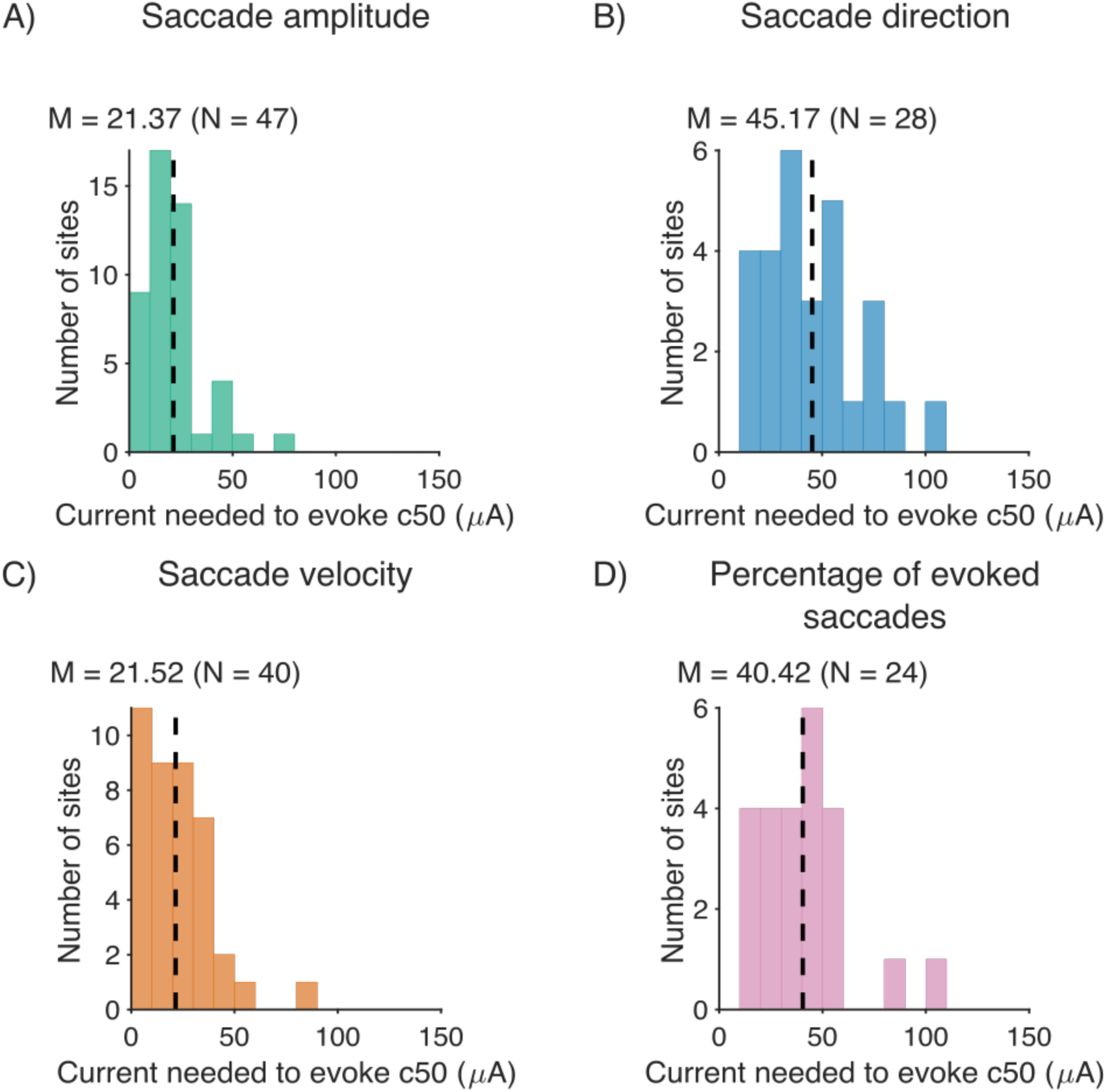
Impact of current intensity on saccades evoked by single-site stimulation in the FEF. (A) Histograms showing the current needed to evoke a 50% change in (A) saccade amplitude, (B) saccade direction, (C) saccade velocity and (D) the percentage of trials with evoked saccades. Black vertical lines represent the mean c50 across sites. Note that the *r^2^* of the Naka-Rushton function had to be greater than 0.7 for a site to be included in the analysis, which explains why there is an unequal number of data points for each histogram.

### Dual-site microstimulation in the FEF with equal current intensities

Next, we performed dual-site microstimulation in the FEF to determine how saccades evoked by single-site stimulation in the FEF are combined to produce a dual-site saccade. In oculomotor regions such as the SC, two main models have been proposed to explain how population activity is read-out by downstream regions: vector summation and vector averaging (Katnani et al. 2012; Lee et al. 1988; van Opstal and Goossens 2008; Van Gisbergen et al. 1987). Although the vector average model is thought to provide a more accurate prediction of saccade amplitude and direction in the SC (Katnani et al. 2012), it is unclear if a similar computation is used to decode population activity in the FEF. Here, we tested the predictions of the vector sum and vector average models against a new polar average model in which amplitude and direction are represented separately by transforming the two constituent saccades into polar coordinates prior to averaging.

The predictions of the three models computing using simulated single-site saccades are shown in Figure 5. With respect to saccade amplitude, the vector sum model overestimated the amplitude of the saccade relative to the polar average model when the difference in theta between the two single-site saccades was relatively small (Figure 5A), whereas the predictions of the two models were similar when the difference in theta was relatively large (Figure 5B). In contrast, the predictions of the vector average and polar average models were similar when the difference in theta between the two single-site saccades was relatively small (Figure 5A), whereas the former underestimated the amplitude of the saccade relative to the latter when the difference in theta was relatively large (Figure 5B). With respect to saccade direction, the predictions of the vector sum and vector average models were always identical. The predictions of the polar average model were also identical to the predictions of the vector sum and vector average models when there was no difference in amplitude between the two single-site saccades (Figure 5C). However, the predictions of the vector sum and vector average models were biased towards the single-site saccade with the largest amplitude when the difference in amplitude between the two single-site saccades was relatively large (Figure 5D). In contrast, the predictions of the polar average model always represented the midpoint between the two-single site saccades irrespective of the difference in amplitude.

**Figure 5.**
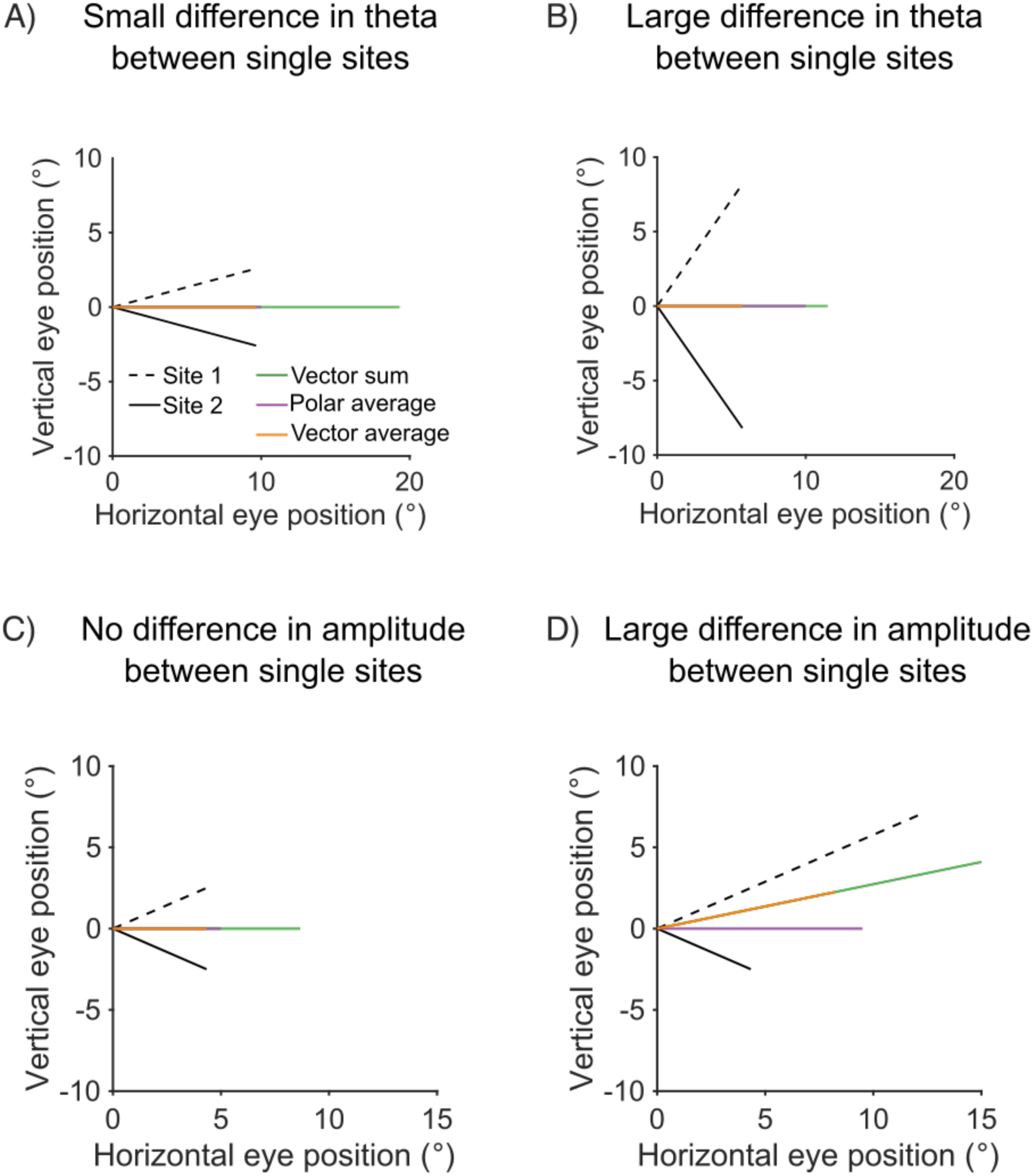
Predictions for the vector sum, vector average and polar average model using simulated single-site saccades. (A) When the difference in theta between the two single-site saccades is relatively small (30°), the vector sum model overestimates the amplitude of the saccade relative to the polar average model, whereas the predictions were more similar for the vector average and polar average models. (B) When the difference in theta between the two single-site saccades is relatively large (110°), the vector average model underestimates the amplitude of the saccade relative to the polar average model, whereas the predictions were more similar for the vector average and polar average models. (C) When there was no difference in amplitude between the two single-site saccades, the predictions for all three models were identical with respect to saccade direction. (D) When the difference in amplitude between the two single-site saccades was relatively large (9°), the predictions for the vector average and vector sum models were pulled towards the saccade with the largest amplitude.

To identify the model that is best able to predict the amplitude and direction of dual-site saccades, the monkeys performed the same task shown in Figure 1A, the only difference being that microstimulation (200 ms, 350 Hz biphasic, 0.25 μS pulse width) was applied simultaneously at two sites along the array with equal current intensities (Figure 6A). The amplitude and direction of saccades evoked by dual-site stimulation was then compared to predictions from two common decoding models (vector sum and vector average) and a new polar average model. Model predictions were computed using the endpoints of the two individual saccades evoked by single-site in the FEF (Figure 2). As can be seen for the example site in Figure 6B, the vector sum model greatly overestimated the amplitude of the dual-site saccade relative to the vector average and polar average computation. In contrast, the vector average model underestimated the amplitude of the dual-site saccade relative to the polar average model. With respect to the direction of the dual-site saccade, the vector sum and vector average models yielded identical estimates that were always biased towards the single-site saccade with the largest amplitude. In contrast, the prediction of the polar average model always coincided with the midpoint between the two single-site saccades.

**Figure 6.**
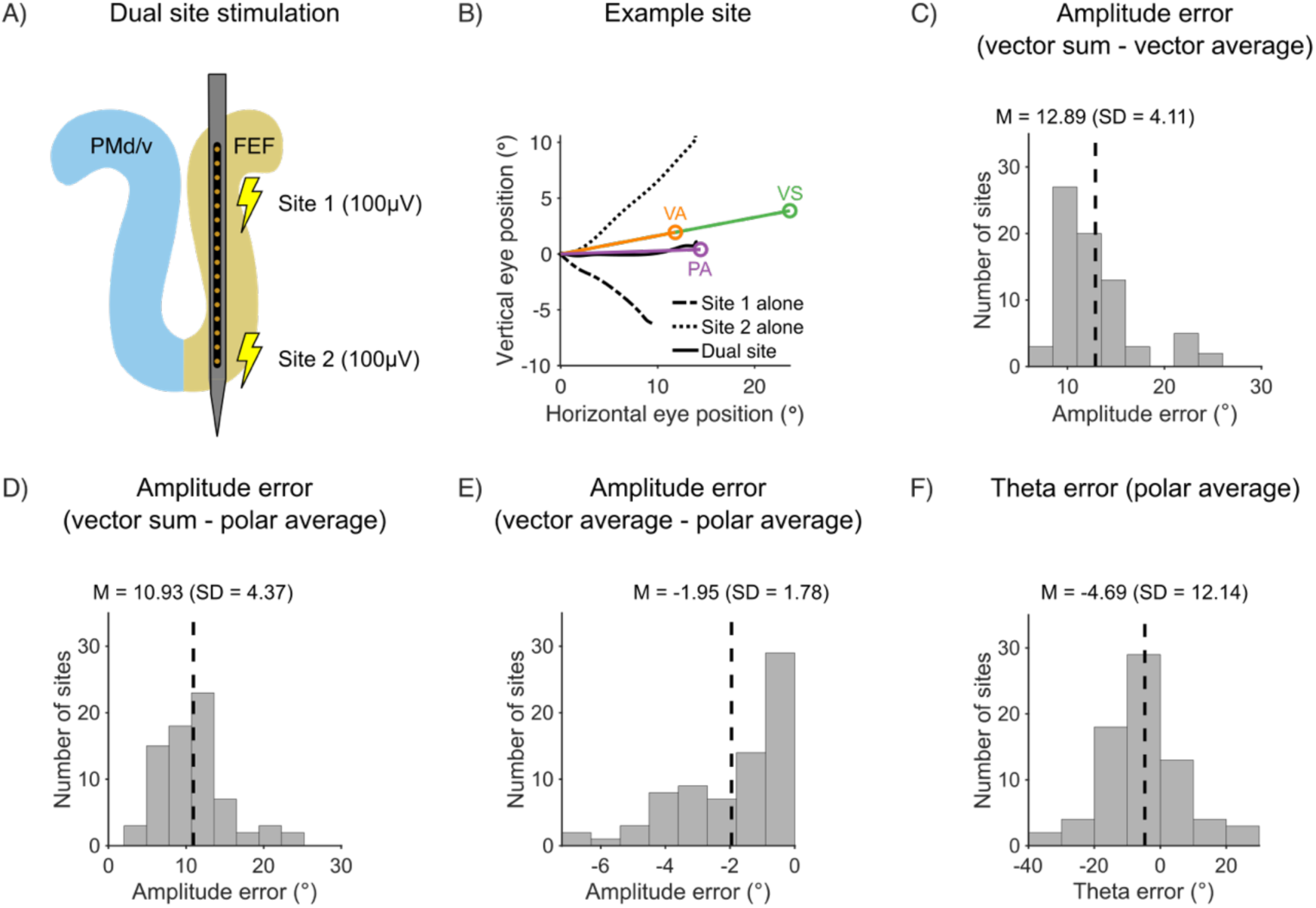
Dual-site FEF microstimulation with equal current intensities. (A) Microstimulation was applied simultaneously at two contacts on the linear microelectrode array. (B) Example site showing predictions for the vector sum, vector average and polar average models. (C) Scatter plot showing amplitude error for the vector sum model plotted against amplitude error for the vector average model. (D) Scatter plot showing amplitude error for the vector sum model plotted against amplitude error for the polar average model. (E) Scatter plot showing amplitude error for the vector average model plotted against amplitude error for the polar average model. (F) Histogram showing theta error for the polar average model and the vector average/sum model.

To explore if a similar pattern was found across all sites, we computed amplitude error by subtracting the amplitude of the dual-site saccade from the amplitude of the model predictions. As such, positive error values indicate an overestimation of the amplitude of the dual-site saccade whereas negative values represent an underestimation of the dual-site saccade. Next, we computed the difference in direction between the dual-site saccade and the model predictions. Given that the predictions for the vector sum and vector average models were always biased towards the larger single-site saccade, whereas those for the polar average model always represented the midpoint (Figure 5D), we computed directional error by first taking the absolute difference in theta between the dual-site saccade and the prediction for the polar average model. The sign of the error was then constrained to be negative when the dual-site saccade was pulled towards the smaller saccade and positive when it was pulled towards the larger saccade. That way, negative error values represent situations in which the polar average model was more accurate than the vector average model at predicting the direction of the dual-site saccade, whereas positive error values represent situations in which the vector sum and vector average models were more accurate than the polar average model at predicting the direction of the dual-site saccade.

Consistent with that was observed across several example sites, paired-sample *t*-tests showed that the vector sum model overestimated the amplitude of the dual-site saccade by 12.89° (*SD* = 4.11°) relative to the vector average model (*t* (72) = 26.82, *p* < 0.001) (Figure 6C). Relative to the polar average model, the vector sum model overestimated the amplitude of the dual site saccade by 10.93° (*SD* = 4.37°) (*t* (72) = 21.37, *p* < 0.001) (Figure 6D). When we compared the vector average model with the polar average model, results showed that the vector average model underestimated the amplitude of the dual-site saccade by 1.95° (*SD* = 1.78°) relative to the polar average model (*t* (72) = -9.36, *p* < 0.001) (Figure 6E). Thus, the polar average model was more accurate at predicting the amplitude of the dual-site saccade than the vector sum and vector average models. With respect to saccade direction, the mean theta error across all sites was -4.69° (*SD* = 12.14°) and a one-sample *t*-test showed that the distribution was significantly different from zero (*t* (72) = -3.3, *p* < 0.001) (Figure 6F). Given that negative error values represent situations in which the polar average model was more accurate at predicting the direction of the dual-site saccades, these findings suggest that there was more directional error for the vector average and vector models than the polar average model.

### Dual-site microstimulation in the SC with equal current intensities

Next, we investigated if the decoding model used to read-out activity in the FEF is the same as that used to read-out activity in the SC. To do so, we leveraged data from a previous study (Katnani and Gandhi 2011) that applied microstimulation to two sites in the SC using single electrodes (Figure 7A). As was the case for the FEF, we compared the amplitude and direction of saccades evoked by dual-site stimulation in the SC to predictions from the vector sum, vector average and polar average models (Figure 7B). Consistent with that was observed in the FEF, paired-sample *t*-tests showed that the vector sum model overestimated the amplitude of the dual-site saccade by 12.89° (*SD* = 6.37°) relative to the vector average model (*t* (18) = 8.82, *p* < 0.001) (Figure 7C). Relative to the polar average model, the vector sum model overestimated the amplitude of the dual-site saccade by 10.18° (*SD* = 6.3°) (*t* (18) = 7.05, *p* < 0.001) (Figure 7D). When we compared the vector average model with the polar average model, results showed that the vector average model underestimated the amplitude of the dual-site saccade by 2.72° (*SD* = 2.16°) relative to the polar average model (*t* (18) = -5.49, *p* < 0.001) (Figure 7E). Thus, the polar average model was more accurate at predicting the amplitude of the dual-site saccade in the SC than the vector sum and vector average models. With respect to saccade direction, results showed that mean theta error across all sites was -2.93 (*SD* = 14.03). However, the distribution did not differ from zero when a one-sample *t*-test was performed (*t* (18) = -0.91, *p* = 0.37) (Figure 7F). Thus, all three models were equally accurate when predicting the direction of saccades evoked by dual-site stimulation in the SC.

**Figure 7.**
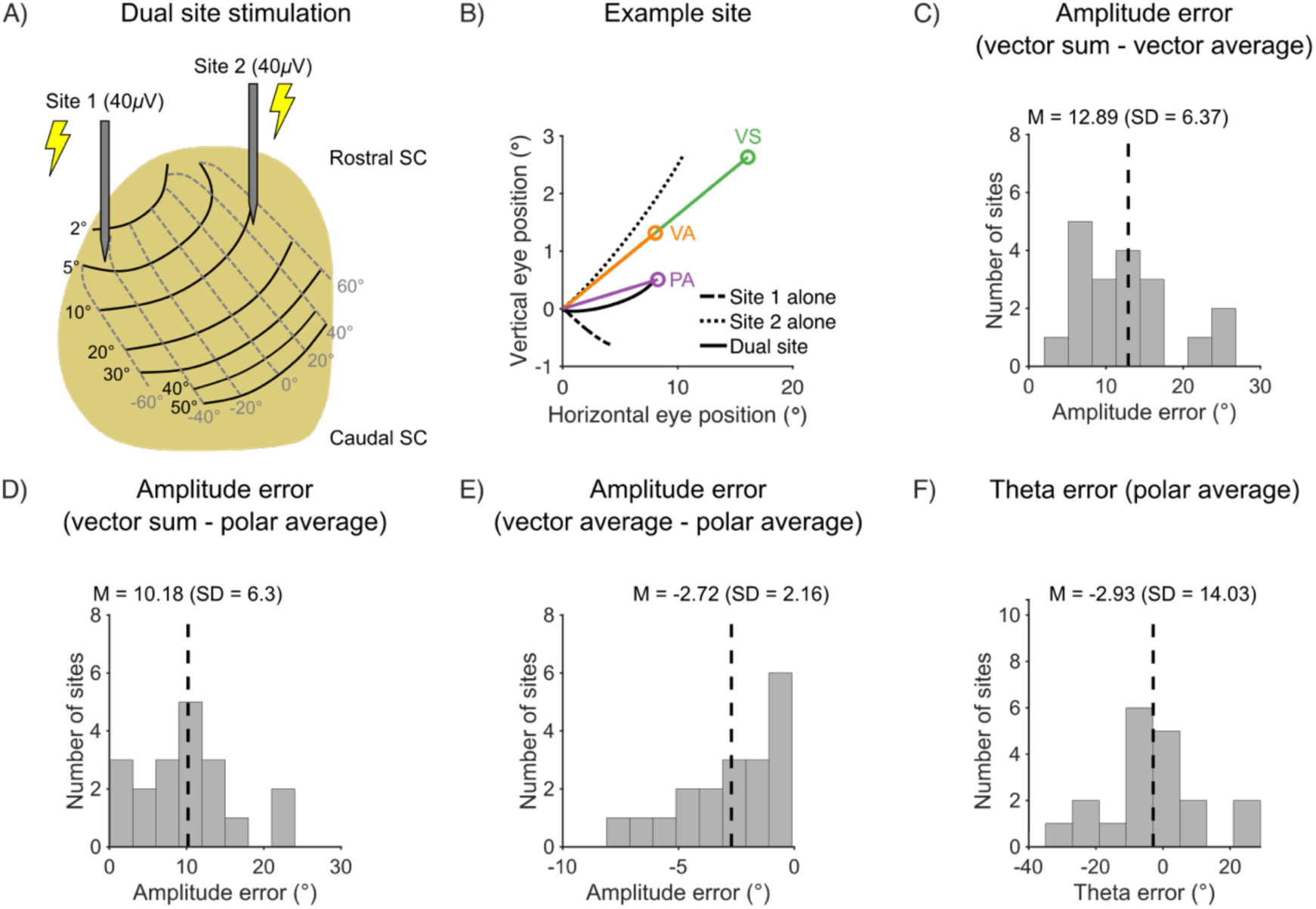
Dual-site SC microstimulation with equal current intensities. (A) Microstimulation was applied at two sites in the SC using single electrodes. (B) An example site showing the predictions of the vector sum, vector average and polar average models. (C) Scatter plot showing amplitude error for the vector sum model plotted against amplitude error for the vector average model. (D) Scatter plot showing amplitude error for the vector sum model plotted against amplitude error for the polar average model. (E) Scatter plot showing amplitude error for the vector average model plotted against amplitude error for the polar average model. (F) Histogram showing theta error for the polar average model and the vector average/sum model.

### Dual-site microstimulation in the FEF with unequal current intensities

Finally, we tested the predictions of the vector average and polar average models when two sites in the FEF were stimulated with unequal current intensities. These two models were significantly more accurate than the vector sum model at predicting the amplitude of saccades evoked by dual-site stimulation with equal current intensities in the FEF (Figure 6) and the SC (Figure 7). For this reason, and the fact that the vector sum and vector average models yield identical estimates of saccade direction (Figure 5D), we limited our focus to the vector average and polar average models for this experiment. The monkeys performed the same task shown in Figure 1A, the only difference being that microstimulation (200 ms, 350 Hz biphasic, 0.25 μS pulse width) was applied simultaneously at two sites along the 16-channel linear microelectrode array with unequal current intensities (Figure 8A). The current intensity at one of the sites on the array was held constant at 30 or 50 μA (termed “Site 1”) whereas the current at the other site (termed “Site 2”) was varied between 15 and 120 μA.

**Figure 8.**
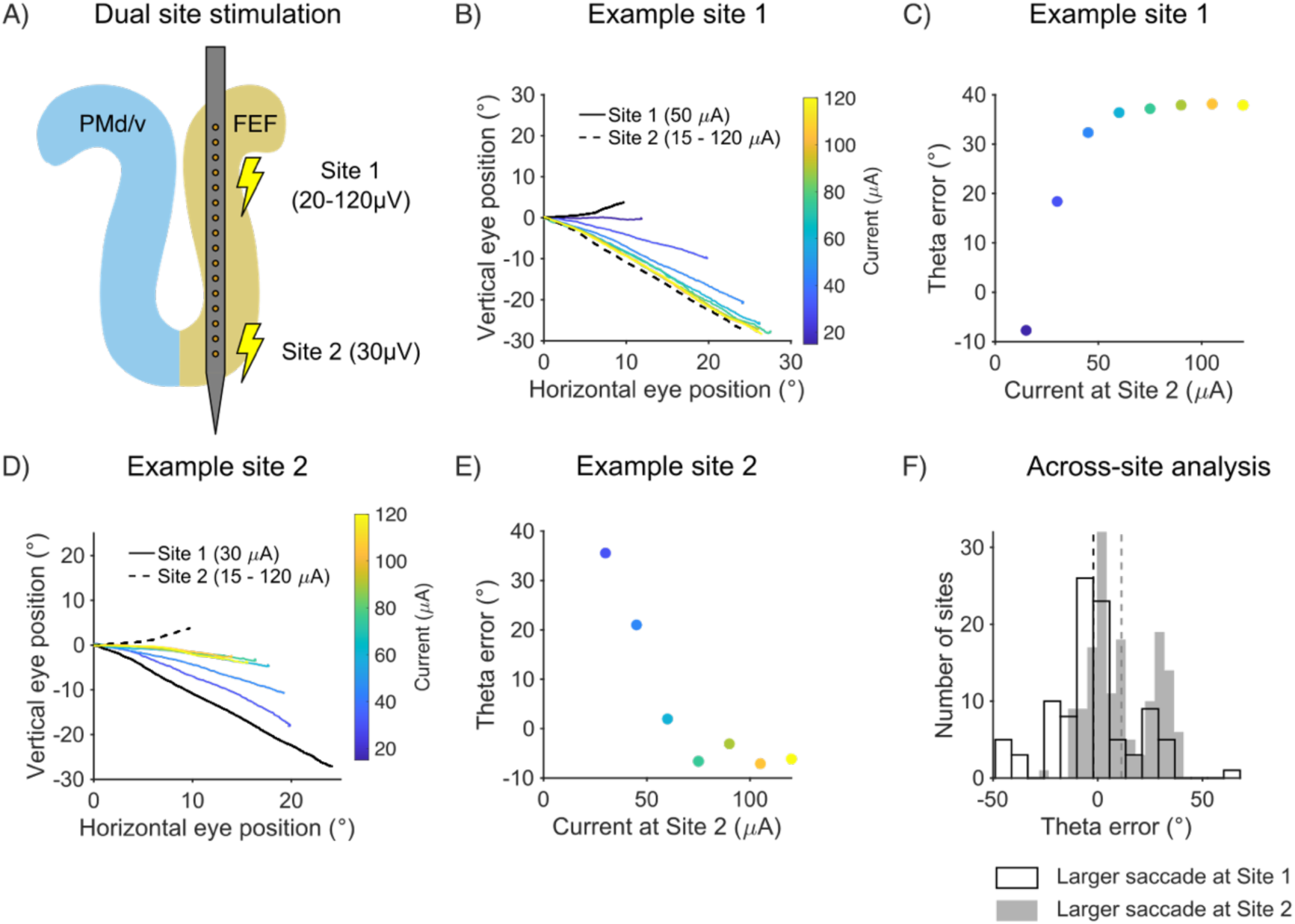
Dual-site FEF microstimulation with unequal current intensities. (A) Microstimulation was applied simultaneously at two contacts on the linear microelectrode array. The current at Site 1 was held constant at 30 or 50 μA, whereas the current at Site 2 was varied between 15 and 120 μA. (B) Dual-site saccades for an example site in which the amplitude of the saccade evoked by stimulating Site 2 alone was larger than the amplitude of the saccade evoked by stimulating Site 1 alone. (C) Theta error for the example site in (B). Negative error values represent situations in which the polar average model was more accurate than the vector average model at predicting the direction of the dual-site saccade and vice versa for positive error values. (D) Dual-site saccades for an example site in which the amplitude of the saccade evoked by stimulating Site 1 alone was larger than the amplitude of the saccade evoked by stimulating Site 2 alone. (E) Theta error for the example site in (D). Negative error values represent situations in which the polar average model was more accurate than the vector average model at predicting the direction of the dual-site saccade and vice versa for positive error values. (E) Histograms showing theta error under two conditions: when the amplitude of the Site 2 saccade was larger than the amplitude of the Site 1 saccade (gray histogram and gray dashed line indicating the mean), and when the amplitude of the Site 1 saccade was larger than the amplitude of the Site 2 saccade (white histogram and black dashed line indicating the mean).

Mean saccade vectors for an example site are shown in Figure 8B. In this case, it is important to note that the amplitude of the saccade evoked by stimulating Site 2 alone was larger than the amplitude of the saccade evoked by stimulating Site 1 alone. Furthermore, as the current at Site 2 increased, the direction of the dual-site saccade shifted towards the saccade that was evoked by stimulating Site 2 alone. Given that the predictions of the vector average model are typically pulled towards the single-site saccade with the largest amplitude (Figure 5D), it is possible that the polar average model may have been more accurate at predicting the direction of the dual-site saccades when current intensity at Site 2 was relatively low, whereas the vector average model may have been more accurate at predicting the direction of the dual-site saccades when current intensity was relatively high. To test this, we compared the direction of saccades evoked by dual-site stimulation with unequal current intensities to predictions from each model. Again, model predictions were computed using the endpoints of the two individual saccades evoked by single-site in the FEF (Figure 2). Theta error was also computed in same manner such that negative error values represent situations in which the polar average model was more accurate than the vector average model at predicting the direction of the dual-site saccade and vice versa for positive error values. As can be seen in Figure 8C, the polar average model was more accurate at predicting the direction of the dual site saccades when current intensity was relatively low (indicated by negative theta error values) whereas the vector average was more accurate at predicting the direction of the dual site saccades when current intensity was relatively high (indicated by positive theta error values). The opposite pattern of results was found for the site shown in Figure 8D. In this case, the amplitude of the saccade evoked by stimulating Site 1 alone was larger than the amplitude of the saccade evoked by stimulating Site 2 alone. Consequently, the vector average model was more accurate at predicting the direction of the dual site saccades when stimulation amplitude was relatively low whereas the polar average model was more accurate at predicting the direction of the dual site saccades when stimulation amplitude was relatively high (Figure 8E).

To explore if a similar pattern was observed across all sites, the data were divided into two separate conditions. Firstly, we identified sites in which the amplitude of the saccade evoked by stimulating Site 2 alone was larger than the amplitude of the saccade evoked by stimulating Site 1 alone. For these sites, we predict that the vector average model will be more accurate at predicting saccade direction. As shown in Figure 8B, the dual-site saccades exhibited shift towards the larger saccade under these conditions. Secondly, we identified sites in which the amplitude of the saccade evoked by stimulating Site 1 alone was larger than the amplitude of the saccade evoked by stimulating Site 2 alone (Figure 8C). For these sites, we predicted that the polar average model will be more accurate at predicting saccade direction. As shown in Figure 8D, the dual site saccades exhibited a shift towards the direction of the smaller saccade under these conditions. Consistent with this hypothesis, the mean theta error across all sites was 11.28° (SD = 15.38°) when the amplitude of the saccade evoked by stimulating Site 2 alone was larger than the amplitude of the saccade evoked by stimulating Site 1 alone (Figure 8F). A one-sample t-test showed that the distribution was significantly different from zero (t (153) = 9.1, p < 0.001) suggesting that the vector average model was more accurate at predicting the direction of the dual-site saccade under these conditions. In contrast, when the amplitude of the saccade evoked by stimulating Site 1 alone was larger than the amplitude of the saccade evoked by stimulating Site 2, the mean theta error across all sites was -2.12° (SD = 20.01°). Although this suggests that the polar average was more accurate at predicting the direction of the dual-site saccade under these conditions, the distribution did not differ from zero when a one-sample *t*-test was performed (*t* (97) = -1.05, *p* = 0.3).

## Discussion

The goal of this study was to investigate the mechanisms underlying saccade generation in the FEF using single- and dual-site microstimulation (Grosbras et al. 2005; Paus 1996; Schall 2015; Tehovnik et al. 2000; Vernet et al. 2014). First, we investigated the influence of current intensity on saccades evoked by single-site stimulation and found that a significantly larger amount of current is needed to evoke a change in saccade direction relative to saccade amplitude. Next, we investigated how population activity in the FEF is read-out by stimulating two sites simultaneously with equal current intensities and found that a new polar average model was more accurate at predicting the amplitude and direction of saccades evoked by dual-site FEF stimulation than the vector sum and vector average models. After comparing the accuracy of the three models in the FEF, we leveraged preexisting dual-site microstimulation data in the SC (Katnani and Gandhi 2011), to assess the generalizability of the polar average model across other oculomotor regions. Although the polar average model was more accurate than the vector sum and vector average models at predicting the amplitude of dual-site saccades in the SC, consistent with what was observed in the FEF, it was no more accurate than the other models at predicting saccade direction. Finally, we compared the accuracy of the three models when two sites in the FEF were stimulated simultaneously with unequal current intensities. Given that the vector sum and vector average models are biased towards the single-site saccade with the larger amplitude, these two models were more accurate at predicting the direction of the dual-site saccades when current intensity was increased at the site with the larger saccade amplitude. In contrast, the polar average model was more accurate at predicting the direction of the dual-site saccades when current intensity was increased at the site with the smaller saccade amplitude.

Previous research investigating the influence of microstimulation parameters on single-site saccades in the FEF has shown that the probability of evoking saccades increases with microstimulation current intensity (Murphey and Maunsell 2008; Robinson and Fuchs 1969; Tehovnik 1996; Tehovnik and Lee 1993; Tehovnik and Sommer 1997). In two monkeys, Murphey and Maunsell (2008) measured the probability of evoking saccades by stimulating at single sites in the FEF and found that the threshold needed to generate a saccade was 19 µA and 23.6 µA for each animal. Here, we investigated the effects of single-site microstimulation on four metrics: saccade amplitude, saccade direction, saccade velocity and the percentage of evoked saccades. For saccade amplitude and saccade velocity, we found that the mean c50 across sites was within the range reported by Murphey and Maunsell (2008). In contrast, the mean c50 for saccade direction was significantly larger than the mean c50 for saccade amplitude suggesting that a greater amount of current is needed bring about systematic changes in saccade direction. Finally, we found that a mean current intensity of 40.42 µA was needed to bring about a 50% decrease from the maximum percentage of evoked saccades. Although this is higher than the probability values reported by Murphey and Maunsell (2008), saccades were evoked during free viewing in their task, whereas in our task monkeys were required to fixate on a central point until 50 ms before stimulation onset. Research has shown that a larger amount of current is needed to evoke saccades in the FEF and SC when fixating on a target, relative to free viewing (Bruce et al. 1985; Goldberg et al. 1986; Sparks and Mays 1983; Tehovnik et al. 1999), which could explain the discrepancy between these results.

Our finding that the c50 for saccade direction was significantly larger than the c50 for saccade amplitude is particularly important and warrants further explanation with respect to the underlying neural mechanisms. The first thing to note is that the c50 for saccade amplitude and saccade direction appears to be similar for the SC. When investigating the impact of microstimulation train duration on saccades evoked by single-site stimulation in the SC, Stanford et al. (1996) found that a similar train duration was needed to bring about a change in saccade amplitude and direction. This contrasts with what we observed in the FEF when microstimulation current intensity was varied. One explanation for this discrepancy stems from the differing topographical organization of the two regions. In the FEF, neurons encoding for saccade amplitude are topographically organized such that small and large saccades can be evoked by stimulating the ventrolateral and dorsomedial portions of the FEF, respectively (Bruce et al. 1985; Dias and Segraves 1999; Robinson and Fuchs 1969). Neurons encoding for saccade direction, on the other hand, are organized more locally in FEF such that progressive changes in the direction of evoked saccades, along with periodic reversals, are observed when an electrode is advanced down the bank of the arcuate sulcus. In the SC, neurons encoding for saccade amplitude and direction are topographically organized and form a continuous map of visual space (Robinson 1972; Schiller and Stryker 1972; Van Opstal et al. 1990). As such, single-site microstimulation of the SC would be expected to activate neighboring cells with similar preferences for saccade amplitude and direction resulting in similar c50 values. Although microstimulation in the FEF would also be expected to activate neurons with similar preferences for saccade amplitude, the preference of these neurons for saccade direction might be more variable resulting in different c50 values for saccade amplitude and direction in this region.

Another important question is how population activity in the FEF and SC is read-out to generate a saccade. To address this in the present study, we compared the amplitude and direction of saccades evoked by dual-site stimulation with equal current intensities to the predictions of two traditional decoding models (e.g., the vector summation and vector averaging), and a new polar average model in which amplitude and direction are represented separately. Results showed that the polar average model was more accurate at predicting the amplitude and direction of dual-site saccades than the vector average and vector sum models. However, when predictions for the three models were computing for preexisting dual-site microstimulation data in the SC (Katnani and Gandhi 2011), we found that although the polar average model was more accurate than the vector average and vector sum models at predicting the amplitude of dual-sites saccades, it was no more accurate at predicting saccade direction. These findings suggest that a different computation may be used to read-out population activity in the FEF and SC, consistent with studies showing that the computation used to decode population activity depends on several factors including the nature of the task (Ferrera 2000; Liu and Wang 2008; Webb et al. 2010; Zohary et al. 1996). In the middle temporal area, prior work has shown that the brain employs a winner-take-all computation when monkeys have to perform a coarse motion discrimination task (Salzman and Newsome 1994) and a vector average computation when they have to pursue a moving target (Groh et al. 1997). Our results extend these findings as they suggest that the brain may use a different computation to read-out population activity even in regions like FEF and SC that have similar functional roles.

Even though the polar average model more accurately predicted the amplitude and direction of saccades evoked by stimulating the FEF with equal current intensities, how such a computation is implemented in the brain remains unclear. As described above, neurons encoding for saccade amplitude and direction in the SC form a continuous map of visual space (Robinson 1972; Schiller and Stryker 1972; Van Opstal et al. 1990) such that a single-site saccade is represented as a Gaussian-shaped mound of activity with the peak of the mound on the map corresponding to the amplitude and direction of the saccade (Gandhi and Katnani 2011; Ottes et al. 1986; Port et al. 2000). When two sites in the SC are stimulated simultaneously with equal current intensities, the two respective activity profiles are combined to form a single mound, which is then read-out by downstream regions as a single saccade command. Given that the FEF has a more heterogeneous topographical organization than the SC, especially with respect to saccade direction, neurons in this region may be better conceptualized as a bank of Gaussian-tuned cells, each sensitive to different saccade amplitudes and directions, rather than a continuous motor map. Downstream regions may then average across this bank of neurons to decode the amplitude and direction of a saccade, similar to how population activity is decoded in the primary visual cortex (Berens et al. 2012; Graf et al. 2011; Kang et al. 2004; Ringach et al. 1997; Stringer et al. 2021). In this respect, the polar average model might be more accurate than the vector average model at predicting the endpoints of saccades evoked by equal-intensity, dual-site FEF stimulation because amplitude and direction are represented separately in this model, unlike vector average model.

An important characteristic of neural population responses is their heterogeneity, with some neurons exhibiting high activity and exerting disproportionate influence, while others exhibit low activity and contribute relatively little to the population code (Chelaru and Dragoi 2008; Stringer et al. 2019; Vinje and Gallant 2000). As a result, it is also important to understand how population activity is decoded under conditions in which there is unequal drive to two groups of neurons, such as when the current is held constant at one site (termed “Site 1”) and systematically varied at another (termed “Site 2”). In the FEF, we observed that the direction of the dual-site saccades was biased away from the site with the lower current intensity and towards the site with the higher current intensity. When the amplitude of the saccade evoked at the site with the higher current insensitivity was larger than that at the site the lower current intensity, the vector sum and average models provided a better prediction of saccade direction than the polar average model. The opposite was true when the amplitude of the saccade evoked at the site with the higher current insensitivity was smaller than that at the site the lower current intensity. However, this effect was generally weaker and did not reach statical significance, possibly because the minimum current used at Site 1 (i.e., 30 µA) was above the threshold needed to evoke a saccade in the FEF, limiting the extent to which the dual-site saccades could be biased towards Site 2. In addition to the task-dependent flexibility described above task (Ferrera 2000; Groh et al. 1997; Liu and Wang 2008; Salzman and Newsome 1994; Webb et al. 2010; Zohary et al. 1996), research has shown neurons with high firing rates and low trial-to-trial variability contribute disproportionately during population decoding (Graf et al. 2011; Zavitz and Price 2019). Our results extend these findings as they suggest that the brain may flexibly combine amplitude and direction information from the FEF in generating saccadic plans.

## Acknowledgements

This work was funded by NIH grant EY022928 and a Research to Prevent Blindness Career Development Award to MAS, NIH P30EY008098, the Eye & Ear Foundation of Pittsburgh, and the Fox Center for Vision Restoration through an Ocular Tissue Engineering and Regenerative Ophthalmology Fellowship to RK. RJ was supported by an unrestricted grant to the University of Pittsburgh Department of Ophthalmology from Research to Prevent Blindness. We are grateful to the staff of the Division of Laboratory Animal Resources at the University of Pittsburgh for animal care.

